# Exploring the impacts of human breast milk functional lipidome on infant health and growth outcomes in early life using lipid bioinformatics

**DOI:** 10.1101/2024.07.25.605136

**Authors:** Moganatharsa Ganeshalingam, Sukhinder Cheema, Carol L. Wagner, Thu Huong Pham, Samantha Enstad, Chloe Andrews, Dmitry Grapov, Flavia Esposito, Sarbattama Sen, Raymond Thomas

**Affiliations:** School of Science and the Environment, Grenfell Campus, Memorial University of Newfoundland, Corner Brook, NL, Canada; Department of Biochemistry, Memorial University of Newfoundland, St. John’s, NL, Canada; Division of Neonatology, Department of Pediatrics, Shawn Jenkins Children’s Hospital, Medical University of South Carolina, United States; Neonatal Intensive Care Unit, Orlando Health Winne Palmer Hospital for Women and Babies, Orlando, FL, United States; Department of Pediatric Newborn Medicine, Brigham and Women’s Hospital, Boston, MA, United States; Creative Data Solution (CDS), Colfax, CA, United States; Department of Mathematics, University of Bari Aldo Moro, Bari, Italy; Pediatrics, Women and Infants Hospital of Rhode Island, Providence, RI, United States; Warren Alpert Medical School, Brown University, United States; Department of Biology, University of Western Ontario, London, Ontario, Canada; Biotron Experimental Climate Change Research Centre, University of Western Ontario, London, Ontario, Canada

**Keywords:** Breast milk lipids, maternal body mass index, bioinformatics tools, Infant growth parameters, Infant Atopic Disease

## Abstract

Human breast milk lipidome is complex, and how changes in the functional lipid metabolism converge systematically to alter infants’ health outcomes is poorly understood. We used human breast milk and infant-mother dyads as a test system to demonstrate how the application of improved lipid bioinformatics can be effective in discerning systematic changes in functional lipid metabolism providing novel discoveries of how lactational programming in early life can influence infant health and growth outcomes. The study consisted of 40 mother-infant dyads where breast milk, maternal diet, infant anthropometrics [fat mass index (FMI), length z score, BMI z score, fat-free mass index (FFMI)], and infant atopic disease outcome (ear infection, cold, wheezing, diarrhea, and eczema) were collected at one and four months postpartum. Integrated Lipid Bioinformatics analyses were conducted using XLSTAT, Metaboanalyst 5.0. R software, Lipid Search, Xcalibur, and Cytoscape software. The results showed breast milk lipidome ordinated into distinct clusters based on maternal BMI status, and differences in developmental and atopic disease outcomes following redundancy analysis. Specifically, lipids from obese mothers clustered with FMI and eczema, while lipids from non-obese mothers clustered with FFM and wheezing. Receiver operating analysis was effective in identifying potential lipid biomarkers that were significantly associated with infant FMI, FFMI, and eczema during early life. Sphingolipid and glycerophospholipid pathways were significantly associated with the altered breast milk lipidome impacting infant development and atopic disease outcome during the first year of life. The findings following the advanced lipid bioinformatics suggest that the breastmilk functional lipid metabolism appears to play a key role in lipid-mediated lactational programming influencing development and atopic disease outcome, and present opportunities for potential dietary intervention in early life.

## 1. Introduction

Lipids play several functional roles in infant growth, gut microbial activity, immunity, and neurodevelopment^1,2^. Breast milk lipids are secreted in the form of fat globules, which consist of a central core of lipids surrounded by a tri-layer membrane called the milk fat globule membrane (MFGM)^3^. The central core of the milk fat globule is rich in triglycerides (TG) with a small amount of monoglycerides (MG), diglycerides (DG), and free fatty acids. The tri-layer membrane is rich in phospholipids and sphingolipids^4^. Several maternal factors including diet, body mass index (BMI), stage of lactation, age, and parity can alter the lipid composition^5^. Studies from this cohort demonstrate that maternal body mass index is a significant modulator of breast milk fatty acids and plasmalogen composition and that the altered lipids appears to be associated with infant growth outcomes, particularly during early life^6,7^. Due to the chemical and functional complexity of breast milk lipids, the determination of the lipidome and associated metabolism can not be achieved by a single analytical and computational approach^8^. Lipidomics refers to a combination of both analytical and computational assessment of the structure and function of the entire composition of the lipids present in a biological sample; and their interaction with other molecular species, as well as ascertaining their biological roles in different metabolic pathways^8,9^. Development of novel and improved analytical techniques especially advances in mass spectrometry techniques, produces a large set of lipid data^10^. The biggest challenge affecting lipidomics when assessing complex samples such as breast milk is explaining the important biological phenomena or features from this large data set^11^. This challenge has necessitated the development of bioinformatics tools that contain improved computational approaches for identifying, quantifying, and further defining metabolic pathways and biomarkers as a resolution^12^. Resolution of these challenges thus allows lipid bioinformatics to help understand the biological meaning inherent in complex data sets. This can be achieved by applying advanced data processing during the identification and quantification of lipids using a series of statistical analyses that include lipid modeling, multivariate, univariate, pathways, and network analyses^11,12^.

Another challenge specifically related to breast milk lipids studies is that the majority of previous studies that focused on assessing the effects of breast milk lipids on infant health outcomes were done using only basic computational approaches. This precludes determining how the complex breast milk lipidome mediates infant health outcome or phenotype from a systems biological approach; and how this may be a feature of lactational programming based on the mother’s physical, physiological, or health status.

Thus, we hypothesized that applying different bioinformatics approaches involving a combination of multivariate, univariate, correlation, pathway, and network analyses would allow us to elucidate improved or novel techniques to assess the breast milk functional lipidome and propose potential associations and mechanisms by which maternal BMI modulates maternal breast milk lipidome, atopic disease, and growth outcomes in infants during early life.

This approach would further serve as a model for other longitudinal studies that link maternal BMI, and breast milk metabolism with infant health, growth, and development. Therefore, our study aims to show the association of maternal body mass index (BMI) on breast milk lipidome, how breast milk lipids could be associated with infant atopic disease and growth outcomes, and the metabolic pathways responsible for the observed phenotype using lipid bioinformatics tools such as multivariate, univariate, correlation, regression, biomarker, pathway, and lipid network analyses.

## 2. Material and Methods

### 2.1. Participants

The current study was a secondary analysis of individuals who consented to participate in a randomized controlled trial of vitamin D supplementation during lactation^13^. A total of 40 of 460 mother-infant dyads were selected from the control group for this trial. The mothers were categorized as non-obese (n=20) ≤30kg/m^2^ and ≥30kg/m^2^ (n=20) based on body mass index (BMI) during pregnancy. Above 90% of infants were exclusively breastfed at one and fourth months postpartum. The criteria restricted formula supplementation to less than 10%. Detailed Inclusion and exclusion criteria for the parent study were reported in a previous publication from the parent study^6,13^.

This study was classified as exempt by the Institutional Review Board of Brigham and Women’s Hospital.

### 2.2. Breast milk collection

Breast milk samples were collected at postpartum visits 1 and 4 months after birth. Breast milk was collected from non-fasting participants using a hospital-grade electric pump. Participants were asked to completely express the opposite breast from which the infant was fed. After the complete expression of the breast, milk was mixed manually and stored at -80°C until further analysis. The processing and storage conditions were previously described^6,7^.

### 2.3. Infant illness data

Data on infant illness (atopic disease) outcomes were collected by study staff through seven months of age during each hospital visit^6,7^. Mothers were asked if their infant had experienced an upper respiratory tract infection, diarrhea, ear infection, eczema, and wheezing during the past month. The data were categorized as dichotomous variables.

### 2.4. Anthropometric measures of infants

Infants underwent whole-body dual-energy X-ray absorptiometry (DXA) scans at 1 and 4 months of age^14^. Global mass, fat mass, and non-obese mass were obtained from DXA results and fat mass and fat-free mass indexes were calculated as previously described^6^. Weight z-scores, length z-scores, and BMI z-scores were calculated from WHO reference data using a 2005 macro (the WHO Child Growth Standards SPSS Syntax File [igrowup. sps]) (Organization & Organization, 2006). The infants’ weight and length were measured during postpartum visits at one and four months and have been described in a previous publication^13^. The measurements were classified as continuous variables during analysis.

### 2.5. Maternal Characteristics

Maternal race and gestational age at delivery were collected during the initial survey, and data on maternal fatty acid intake were collected as described in our previous publication^7^.

### 2.6. Extraction and Analysis of complex breast milk functional lipids

Breast milk samples were extracted using the modified Bligh and Dyer method, where 100 µl of breast milk was placed into glass vials. Aliquots of 20µl of 1mg/1ml internal standard (SPLASH mix from Avanti Polar Lipids, Birmingham, AL catalog number 330707) were added and 1 mL methanol (0.01% butylated hydroxytoluene) followed by 1.0 ml chloroform. The sample was homogenized using a vortex for 1 minute, after which 0.8 ml water was added and the sample re-vortexed. The breast milk sample mixture was next centrifuged at 1500 rpm for 10 minutes, and the organic phase was transferred to pre-weighed vials, and then dried under N_2_ gas (Bligh and Dyer.,1959). Methanol: chloroform (1:1 v/v) was added to the recovered lipids to resuspend the breast milk lipids on a mg/mL basis. The lipids extracted were analyzed using C30 reverse-phase liquid chromatography (C30 RPLC) coupled with heated electrospray ionization and high-resolution accurate mass tandem mass spectrometry (UHPLC-C30RP-HESI-HRAM-MS/MS). The Accucore C30 reverse-phase column (150 × 2 mm I.D., particle size: 2.6 μm, pore diameter: 150 Å) used was obtained from Thermo Fisher Scientific (ON, Canada). The mobile phase system consisted of solvent A (acetonitrile: H_2_O 60:40 v/v) and solvent B (isopropanol: acetonitrile: water 90:10:1 v/v/v/v/v) both containing 10 mM ammonium formate and 0.1% formic acid. C30-RPLC separation was carried out at 30 °C (column oven temperature) with a flow rate of 0.2 mL/min, and 10 μL of the lipid extract suspended in chloroform: methanol (1:1 v/v) was injected into the column. The following system gradient was used for separating the lipid classes and molecular species: 30% solvent B for 3 mins; then solvent B increased to 43% over 5 min, then to 50% B in 1 min, then to 90% B over 9 mins, then to 99% B over 8 mins and finally kept at 99% B for 4 mins. The column was re-equilibrated to starting conditions (using 70% solvent A) for 5 minutes before each new injection. Lipid analyses were carried out using a Q-Exactive Orbitrap mass spectrometer controlled by X-Calibur software 4.0 (Thermo Scientific, MO, USA) and coupled with an automated Dionex Ultimate 3000 UHPLC system controlled by Chromeleon software. The following parameters were used for the Q-Exactive mass spectrometer-sheath gas: 40, auxiliary gas: 2, ion spray voltage: 3.2 kV, capillary temperature: 300 °C; S-lens RF: 30 V; mass range: 200–2000 m/z; full scan mode at a resolution of 70,000 m/z; top-20 data-dependent MS/MS at a resolution of 35,000 m/z and collision energy of 35 (arbitrary unit); isolation window: 1 m/z; automatic gain control target: 1e5. The instrument was externally calibrated to 1 ppm using ESI negative and positive calibration solutions (Thermo Scientific, MO, USA). Tune parameters were optimized using a mixture of lipid standards (Avanti Polar Lipids, Alabama, USA) in both negative and positive ion modes^15^.

### 2.7. Identification and quantification of breast milk lipid molecular species

Lipid Search 4.2 (Mitsui Knowledge Industry, Tokyo, Japan) and Xcalibur 4.0 (Thermo Fisher Scientific, Mississauga, ON, Canada) software packages were used for the identification and quantification of lipids. Lipid search identified the lipidome of the samples, Xcalibur was used for manual curation to confirm accurate identification and for qualification of the identified lipids per classes in the samples against the splash standard mix injected.

### 2.8. Statistical analysis

All Statistical analysis was carried out using XLSTAT (2021 Premium version, Addinsoft, Paris, France), Cytoscape, R studio, and Metaboanalyst 5. Outliers were checked using the Grubs test, and data were standardized using Pareto scaling. The normality of the data was checked with the Shapiro-Wilk test. The total breast milk data were divided into membrane lipids and storage lipids based on nomenclature for the milk fat globule composition for further statistical analysis^16^. The workflow of methodology described in figure 1.

**Figure 1.**
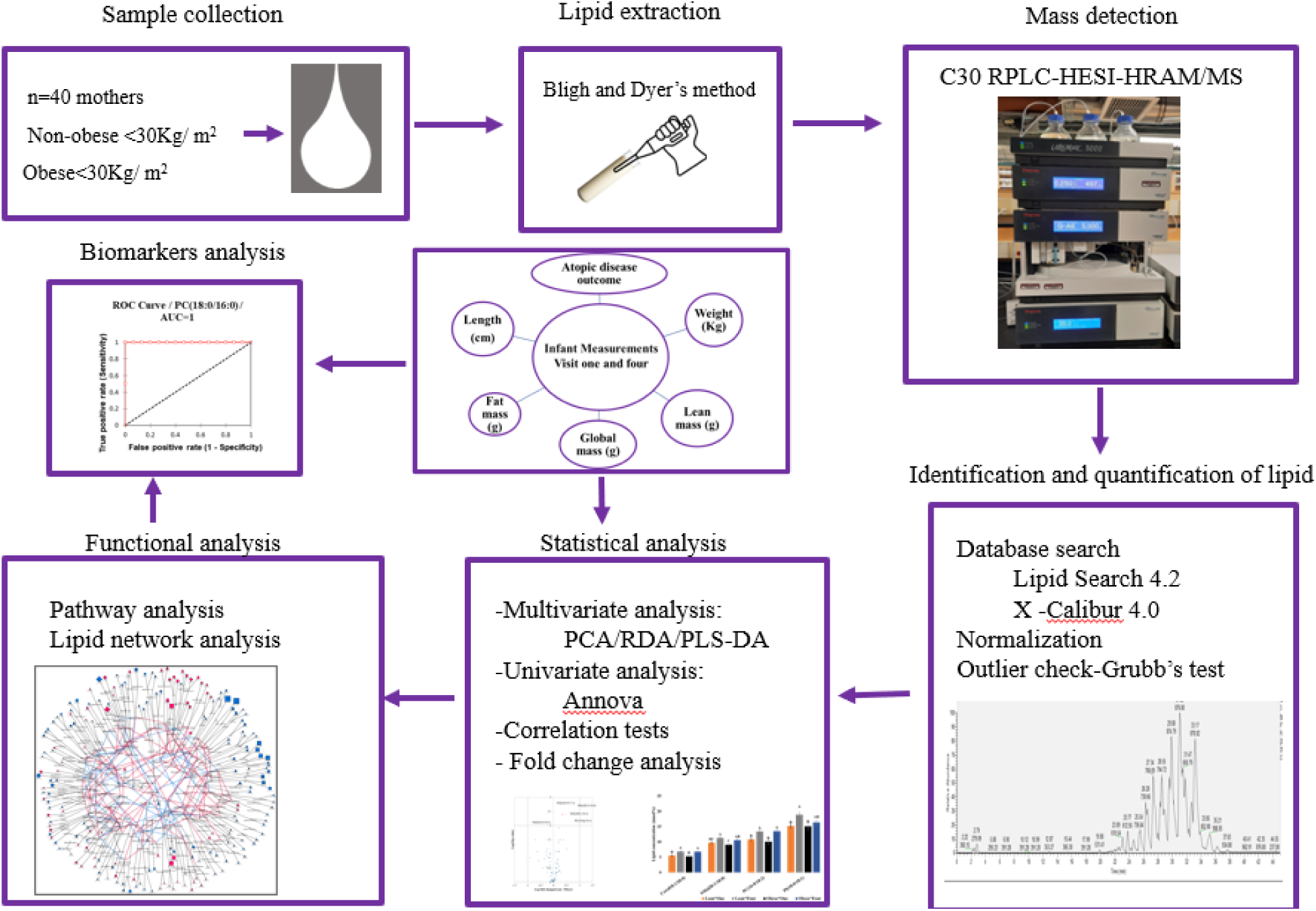
The lipid bioinformatics workflow used in this study.

#### 2.8.1 Application of different Multivariate analysis

Differential analysis (DA), redundancy analysis (RDA), Principal Component Analysis (PCA), and Partial Least Square Discriminant Analysis (PLS-DA) multivariate approaches were applied to the data set to determine the best methods to discern the unique or important functional lipid clusters based on maternal status and time post-partum ( one and four months) milk was collected, as well as to find significant correlations and patterns between breast milk lipids, maternal groups, time post-partum (visit one and visit four), and infant clinical parameters. The volcano plots were created following differential analysis to identify differentially expressed lipids according to two factors: lipids and maternal status (two levels: non-obese and obese) with a p-value of 0.05 and threshold set at (0.01), and the Benjamin-Hochberg method used for post hoc correction.

#### 2.8.2 Student’s T-test

The student’s t-test was used to compare the means of membrane lipids and storage lipids of non-obese and obese mothers’ breast milk. The lipids clustered with the obese and non-obese maternal status ordinated in each quadrant of the RDA biplot were extracted and a student’s t-test was performed to assess the groups’ significance. The results were presented as mean ± standard error and p < 0.05 was considered significant.

#### 2.8.3 Two-way ANOVA

Two-way ANOVA was used to determine the significant lipid molecular species that accounted for the segregation of the groups in different quadrants of the RDA biplots. It was carried out to compare the main effects of maternal status (non-obese and obese), and time post-partum (one and four months).

Two-way ANOVA followed by post hoc Turkey’s test was used to separate the means when the groups were observed to differ from each other at p<0.05 significantly. The results were presented as mean ± standard error.

#### 2.8.4. Correlation test

The Pearson correlation coefficient was computed for each quadrant of the multivariate analysis (RDA) to find the relationship between lipids and infant growth parameters (continuous variable), and the biserial correlation coefficient was computed to find the association between maternal breast milk lipids and infants atopic disease outcome (dichotomous variable) during the first year of life. An ANCOVA regression model was used to adjust covariates such as maternal dietary fat intake, maternal race, maternal age, and maternal education level, on the lipid concentration or levels, and p-value < 0.05 was considered significant.

#### 2.8.5. Biomarker analysis

The receiver operating curve was used to determine the potential biomarkers from the group of lipids that were significantly correlated with clinical outcomes. The area under the curve (AUC) was set to 0.5, and the significance level alpha threshold was set at 0.05.

#### 2.8.6. Lipid-Lipid Network Analysis and Regularized Correlation Network Analysis and Network Mapping

Lipid-lipid partial correlation network analysis was conducted to display the differences present in obese relative to the nonobese cohorts at one and four months postpartum.

Specifically, the changes in lipids observed to be significantly different following RDA and ANOVA analyses were used to generate the lipid-lipid network reflecting the changes present in the lipidome of the non-obese visit (LV) as well as the obese visit (OV) cohorts at one and four-month postpartum. Networks were calculated using the R programming language ^17^ and visualized in Cytoscape ^18^.

Correlations between lipids were calculated based on the high-dimensional undirected graph estimation (huge) method^19^. Relationships between lipids were determined based on Meinshausen-Buhlmann graph estimation and regularization was applied using a stability approach to regularized selections (stars) with lambda specified at 0.21. The direction and magnitude of lipid-lipid associations were expressed as false discovery rate-adjusted^20^. Mapped networks were created to visualize changes in lipids between non-obese and obese cohorts at time post-partum (one and four months)^20^.

Lipid-to-lipid connections were displayed based on identified regularized correlations. The magnitude and direction (positive, negative) of the relationship were determined based on false discovery rate adjusted Spearman correlation, with p-values < 0.05 (FDR)^21^. The network displays significant Spearman correlations among lipids (pFDR < 0.05) as edge colors (positive-red; negative-blue).

Statistical contrasts (p-value < 0.05) in obese relative to non-obese cohorts at one and four months are connected to lipid nodes (dark gray edges). Obese vs non-obese comparisons, which are not statistically different between time points are connected with light gray edges. The magnitude and direction of fold changes in obese relative to non-obese are shown as node size and color (red, increased; blue, decreased). Comparisons at time post-partum (one and four months) are denoted by node border thickness (thin, one month; thick, four months). Lipid classes are denoted with node shape.

### 2.9. Pathway analysis

Pathway analysis was done using Metaboanalyst 5 to reveal the functional impact of maternal BMI on breast milk lipid metabolism. A pathway scatter plot was created according to the p values from the pathway enrichment analysis, and pathway impact values were determined after the pathway topology analysis. KEGG pathway analysis showed the lipids interaction, reaction, and metabolism.

## 3. Results

### 3.1. Demographics of study participants

The demographics of the study participants have already been published ^6,7^. Descriptive characteristics of the mothers and their infants’ dyads are outlined in Table S1. The mean BMI of nonobese mothers was 25.02 kg/m2, while obese mothers were 33.92 kg/m2. The average birth weight mean (±SD) was 3372.55(442.358) g for infants of nonobese mothers, and 3242.80(492.776) g for infants of obese mothers. The mothers were predominantly Caucasian, with a higher proportion of male infants.

### 3.2. Multivariate Statistical Analysis of Lipids

PCA, PLS-DA, and RDA were run to assess the data structure, reduce the data dimensionality, and cluster breast milk lipid molecular species according to maternal status and time postpartum for breast milk collection. The PCA plot of membrane and storage lipids explains 46.46% and 41.24 % of the total variance captured by the first two principal components. PCA did not demonstrate the clustering trends based on maternal group and time postpartum and was not appropriate for the structure of this data set. Similarly, for PLS-DA, the cumulated Q^2^ (predictive ability) index and R^2^Y (coefficient determination) of the membrane and storage lipids were 0.218 and 0.075; and 0.305 and 0.385, which is less than 0.6 representing the set threshold recognized as appropriate for PLSDA models (Figure S1). Thus, the PLS-DA model was unsuitable for this data set, as a Q^2^ and R^2^Y threshold above 0.6 generally indicates good model quality.

In comparison, the RDA output of membrane and storage lipids explained 76.71 % and 60.24 % of the total variance covered by the first two components (Figure 2). The test was statistically significant (p<0.0001). Based on the highly explained variance and visualization trends, we selected RDA as the suitable multivariate analysis for assessing the breast milk functional lipids data set. This multivariate analysis clearly distinguished maternal status with time post-partum (one and four months) based on breast milk lipids.

**Figure 2:**
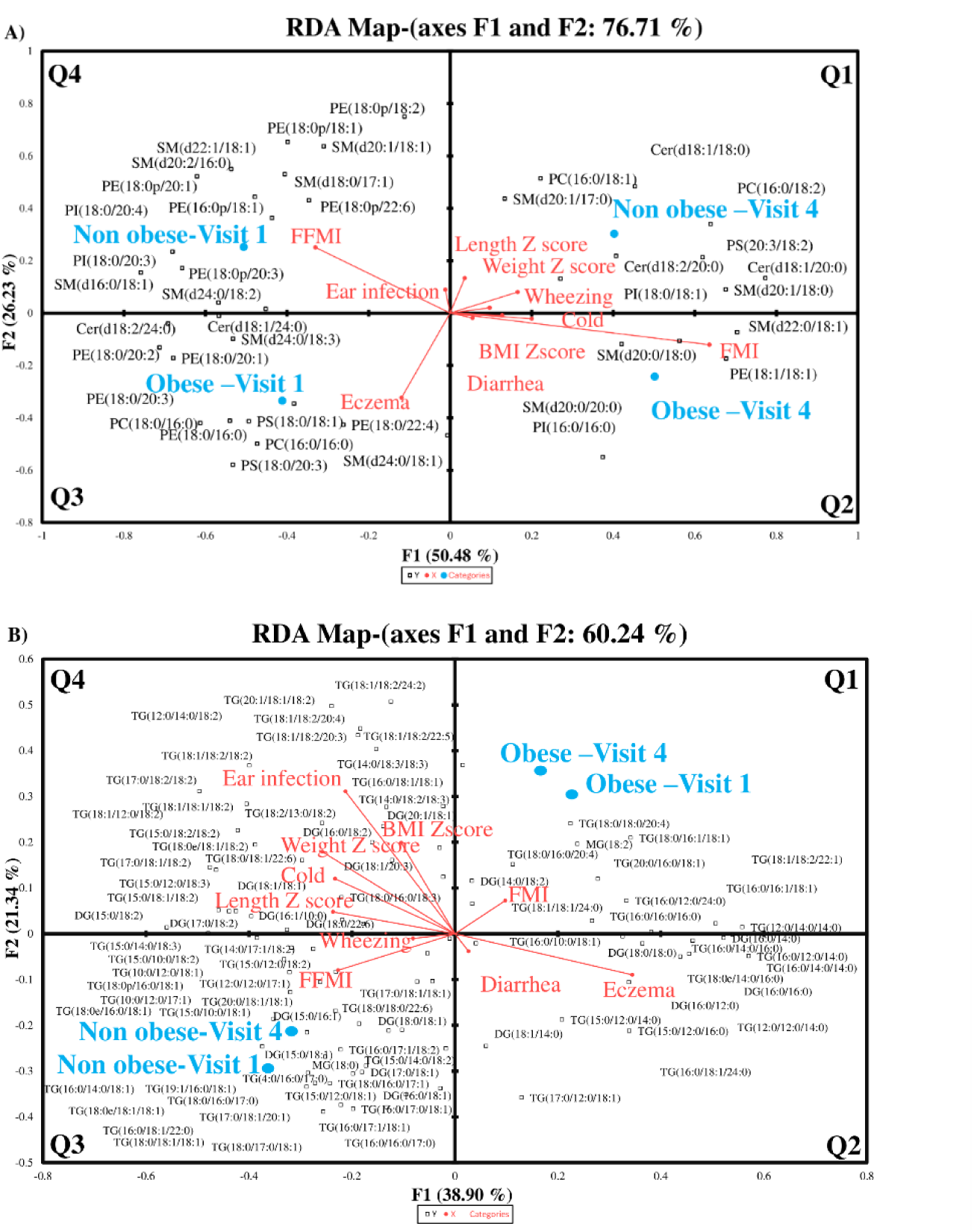
Redundancy analysis (RDA) biplots showing the ordination of breastmilk membrane (A) and storage lipid species (B) based on maternal status, atopic disease, and infant growth outcomes during early life. SM-Sphingomyelin, Cer-Ceramide, PC-Phosphatidylcholine, PE-Phosphatidylethanolamine, PS-Phosphatidylserine, PI-Phosphatidylinositol, TG-Triglycerides, MG-Monoglycerides, and DG-Diglycerides, FMI-Fat Mass Index, FFMI-Fat Free Mass Index Visit 1and Visit 4-one-and four-month post-partum visits

Figure 1 illustrates the ordination of breast milk lipids and their associations with maternal status, time post-partum (one and four months), and various infant clinical parameters. For membrane lipids (Figure 2A), lipid molecular species clustered distinctly into four quadrants based on maternal status and postpartum visits: obese visit 1, non-obese visit 1, obese visit 4, and non-obese visit 4. Quadrants 2 and 4 also showed clustering based on maternal status alone (obese and non-obese). Specific associations were noted with infant health parameters. For example, lipids in Quadrant 1 were associated with length Z score, weight Z score, and wheezing. Quadrant 2 lipids were linked to cold, diarrhea, BMI Z score, and Fat Mass Index (FMI). Quadrant 3 lipids showed a strong association with eczema; and Quadrant 4 lipids were associated with ear infection and Fat-Free Mass Index (FFMI). For storage lipids (Figure 2 B), the variance was primarily attributed to maternal status, with lipid molecular species clustering in quadrants 2 and 4 based on maternal status and postpartum visits. Additionally, Quadrant 1 lipids were linked to FMI, and Quadrant 3 lipids were associated with FFMI and wheezing. These findings underscore the significant impact of maternal status and postpartum visits on the distribution of breast milk lipids and their associations with various infant growth and health outcomes in the first year of life.

### 3.3. Assessment of discriminating variables

Volcano plots (Figure 3) identified the most discriminating lipids based on statistical significance (p-value) and magnitude of change (fold change). The lipids above the threshold limit (horizontal dashed line) were considered biological and statistically significant metabolites. SM (d18:0/17:1), SM (d20:1/18:1), PE (18:0p/18:1), and SM (d20:2/16:0) were upregulated in nonobese mothers’ breast milk, while SM (d24:0/18:1) and TG (18:0/16:1/18:1) were upregulated in obese mothers’ breast milk.

**Figure 3:**
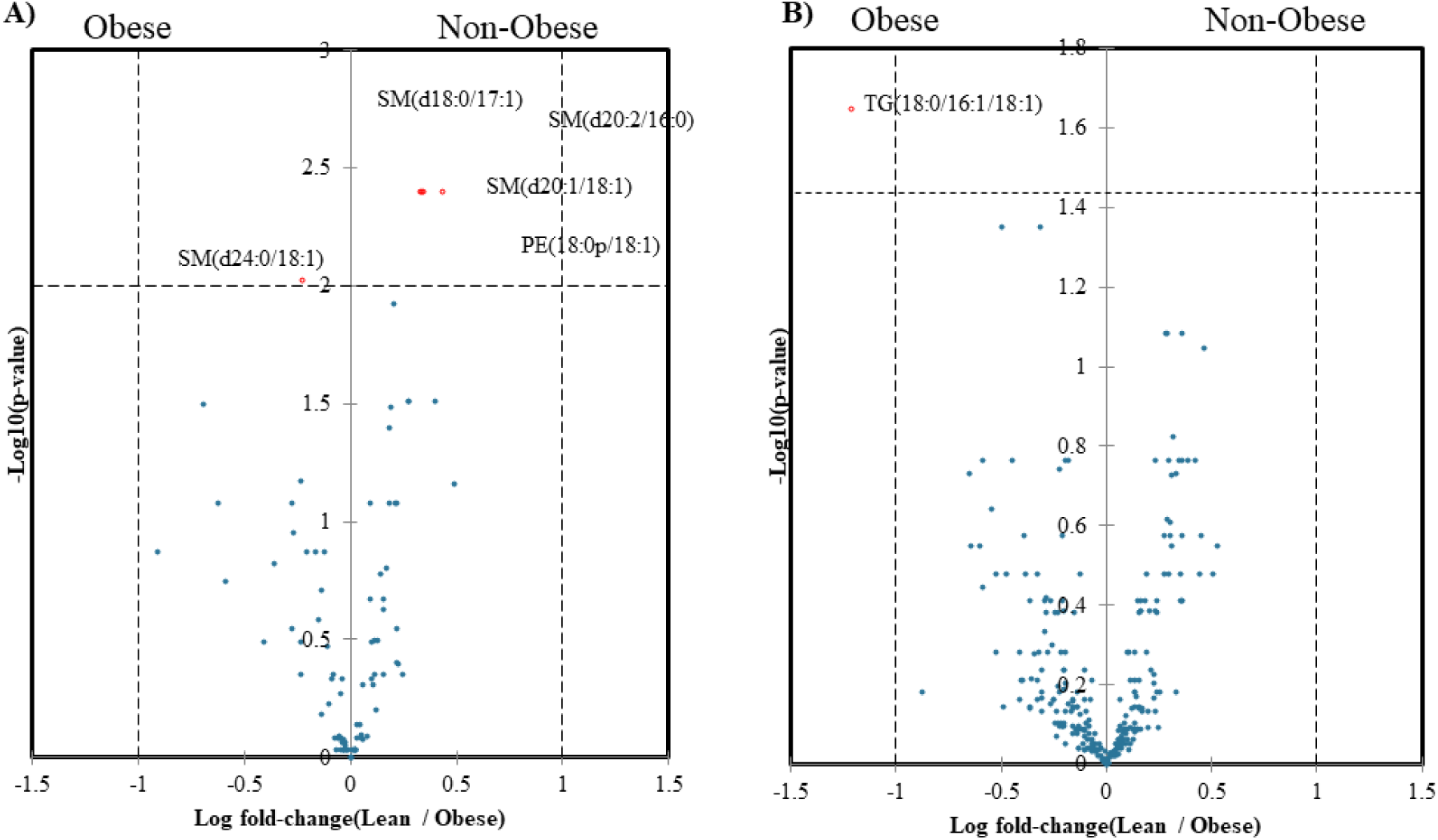
Volcano plot of membrane (A) and storage lipids (B) displaying the comparison between non-obese and obese mothers. Significance threshold p<0.05. Thresh hold (0.01) was set and Benjamini-Hochberg was used for post hoc correction.

### 3.4. Effects of Maternal Body Mass Index and Time Postpartum on the Membrane and Storage Lipids Composition of Breast Milk

The significant breast milk lipids in obese and nonobese mothers’ breastmilk are listed in Table 1. SM, TG, and PI containing saturated fatty acids and TG containing long-chain fatty acids long-chain fatty acids were significantly higher in obese mothers’ breast milk. The sn-2 oleic acid enriched in SM, DG, and MG; short-chain fatty acids enriched TG and MG, and plasmalogen containing PE molecular species were significantly higher in the non-obese mothers’ breast milk. The membrane lipids clustered in quadrants one and two (Fig 4 A and B) were significantly higher in the breast milk of non-obese and obese mothers at the fourth-month post-partum visit. PI (16:0/16:0) and PC (16:0/18:1) were significantly different between obese and non-obese mothers at one-and four-month post-partum visits, respectively. The membrane lipids clustered in quadrants three and four (Fig 4 C and D) were significantly higher in breast milk of non-obese and obese mothers at one-month post-partum visit. SM (d20:2/16:0), PE (18:0p/18:1), and SM (d20:1/18:1) showed a significant difference between obese and non-obese mothers at one-and four-month post-partum visits. The storage lipid in quadrant one did not show any significant changes with maternal status and visits. TG (16:0/10:0/18:1) and TG (16:0/18:1/18:1) showed significant differences between obese and non-obese at one-and four-month post-partum visits, respectively (Figure 5).

**Figure 4:**
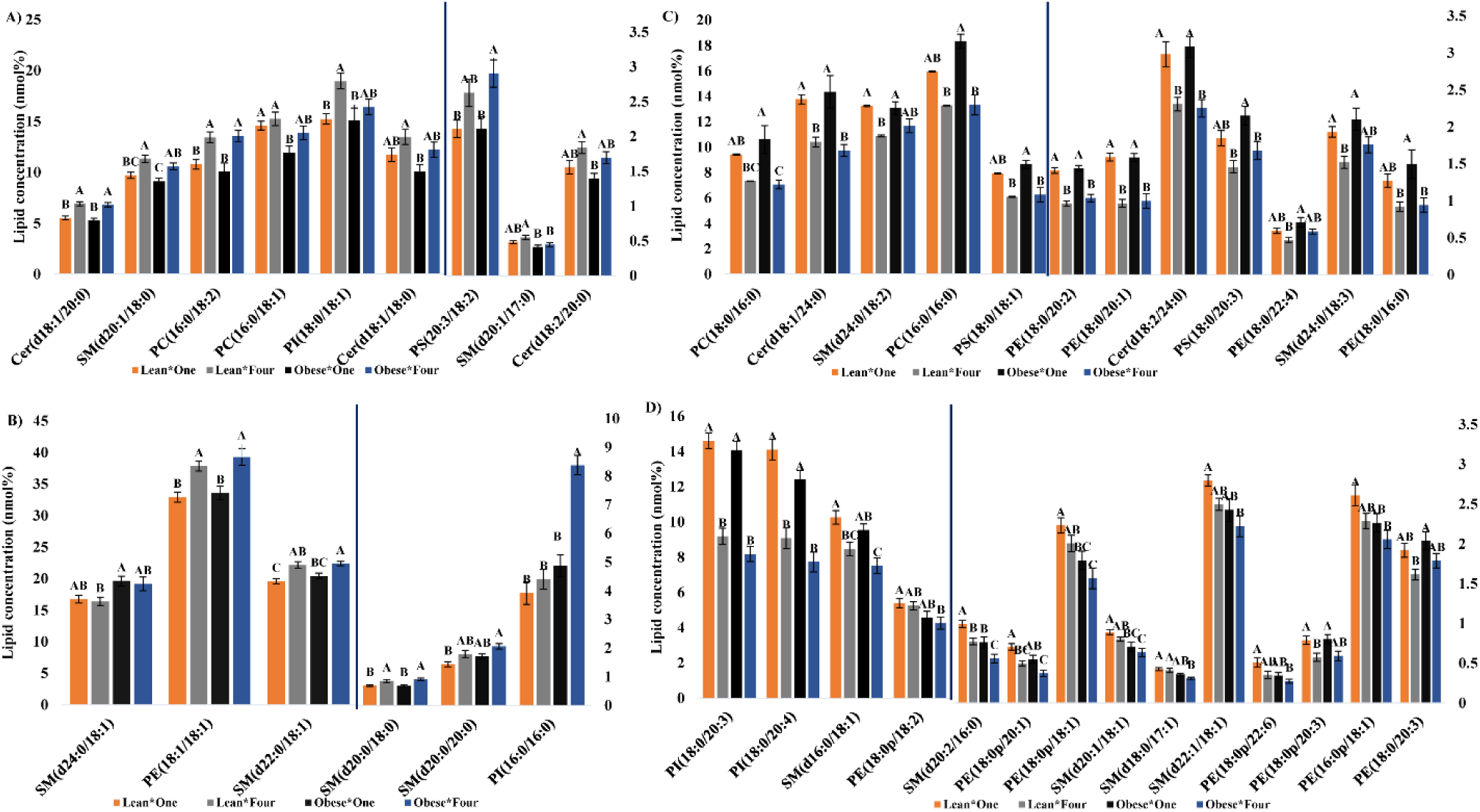
Univariate analysis post-RDA showing the significance of breast milk membrane lipids ordinated in different quadrants of the biplot. Data were analyzed by Two-way ANOVA based on RDA quadrant ordination. Bar charts revealed significant (p<0.05) changes in membrane lipid molecular species in Quadrant 1(A), Quadrant 2(B), Quadrant (C), and Quadrant 4 (D). Data are expressed as nmol% of the total lipids; values are expressed as mean ± SE and different superscripts (a, b, c) are used to denote significant differences between the maternal status and visit. LV1-Non-obese Mother Visit One, OV1-Obese Mother Visit One, LV4-Non-obese Mother Visit Four, OV4-Obese Mother Visit Four. SM-Sphingomyelin, Cer-Ceramide, PC-Phosphatidylcholine, PE-Phosphatidylethanolamine, PS-Phosphatidylserine, PI-Phosphatidylinositol, TG-Triglycerides, MG-Monoglycerides, and DG-Diglycerides.

**Figure 5:**
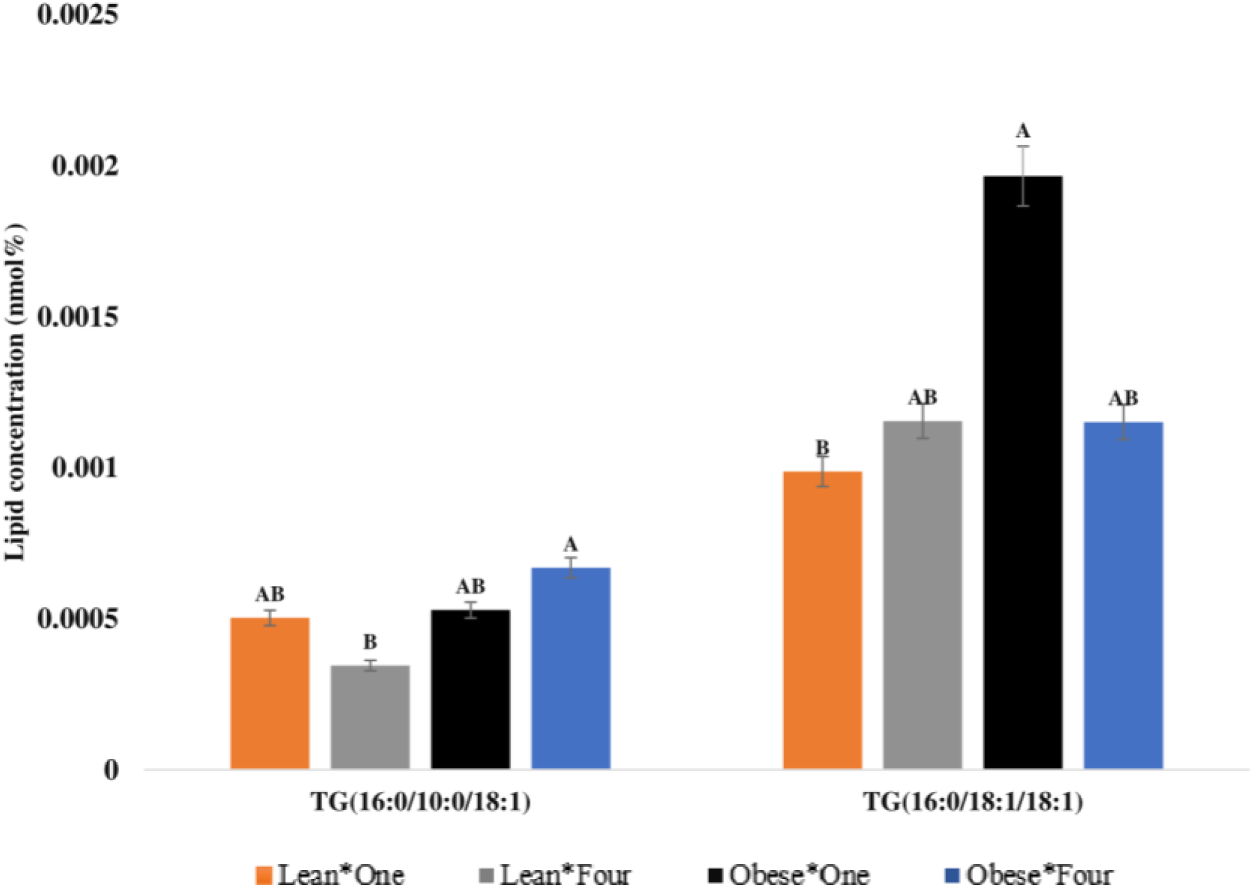
Univariate analysis post-RDA showing the significance of breast milk storage lipids ordinated in different quadrants of the biplot. Bar charts revealed significant (p<0.05) changes in glycerolipid molecular species in quadrant 3. Data are expressed as nmol% of the total lipids; values are expressed as mean ± SE and different superscripts (a, b, c) are used to denote significant differences between the maternal staus and visit. LV1-Non-obese Mother Visit One, OV1-Obese Mother Visit One, LV4-Non-obese Mother Visit Four, OV4-ObeseMother Visit Four. SM-Sphingomyelin, Cer-Ceramide, PC-Phosphatidylcholine, PE-Phosphatidylethanolamine, PS-Phosphatidylserine, PI-Phosphatidylinositol, TG-Triglycerides, MG-Monoglycerides, and DG-Diglycerides.

**Table 1.** Altered Lipids observed in breast milk from non-obese and obese mothers.

| Lipid | Non-obese | Obese | p-values |
| --- | --- | --- | --- |
| SM (d16:0/18:1) | 9.37±0.3 | 8.47±0.33 | 0.049 |
| SM (d20:1/18:1) | 0.84±0.02 | 0.67±0.04 | <0.0001 |
| SM (d22:1/18:1) | 2.65±0.06 | 2.32±0.1 | 0.004 |
| PE (18:0p/22:6) | 0.43±0.04 | 0.31±0.02 | 0.01 |
| PE (16:0p/18:1) | 2.45±0.08 | 2.16±0.08 | 0.014 |
| PE (18:0p/18:2) | 5.32±0.18 | 4.41±0.24 | 0.003 |
| PE (18:0p/20:1) | 0.60±0.03 | 0.46±0.04 | 0.003 |
| PE (18:0p/18:1) | 2.12±0.07 | 1.67±0.09 | 0 |
| SM (d20:2/16:0) | 0.88±0.03 | 0.65±0.05 | 0 |
| SM (d20:0/20:0) | 1.61±0.07 | 1.90±0.08 | 0.009 |
| PI (16:0/16:0) | 4.16±0.84 | 6.72±0.26 | 0.003 |
| TG (18:0e/18:1/18:1) | 0.021±0.002 | 0.015±0.001 | 0.002 |
| TG (18:0/18:1/18:1) | 2.782±0.144 | 2.307±0.111 | 0.002 |
| TG (16:0/17:0/18:1) | 0.119±0.011 | 0.09±0.006 | 0.005 |
| TG (18:0/17:0/18:1) | 0.051±0.005 | 0.037±0.003 | 0.006 |
| TG (18:0/18:0/22:6) | 0.012±0.002 | 0.008±0.001 | 0.027 |
| TG (18:0e/16:0/18:1) | 0.053±0.004 | 0.042±0.003 | 0.013 |
| TG (4:0/16:0/17:0) | 0.005±0.001 | 0.003±0 | 0.031 |
| TG (20:0/18:1/18:1) | 0.474±0.042 | 0.386±0.026 | 0.028 |
| MG (18:0) | 17.046±1.454 | 13.321±0.806 | 0.006 |
| DG (17:0/18:1) | 0.407±0.035 | 0.325±0.021 | 0.012 |
| DG (15:0/18:1) | 0.538±0.04 | 0.441±0.029 | 0.018 |
| DG (18:0/18:1) | 2.986±0.247 | 2.515±0.152 | 0.044 |
| TG (18:0/16:1/18:1) | 0.614±0.027 | 0.282±0.084 | 0 |
| TG (18:0/18:0/20:4) | 0.026±0.001 | 0.019±0.002 | 0 |
| TG (16:0/10:0/18:1) | 0.513±0.019 | 0.42±0.020 | 0.001 |
| TG (16:0/12:0/14:0) | 0.452±0.018 | 0.397±0.018 | 0.034 |
| TG (18:0/16:0/20:4) | 0.067±0.004 | 0.052±0.006 | 0.037 |
| DG (14:0/18:2) | 4.005±0.014 | 3.501±0.02 | 0.04 |
| TG (20:0/16:0/18:1) | 0.223±0.017 | 0.149±0.029 | 0.024 |
| TG (18:1/18:2/20:3) | 0.035±0.003 | 0.023±0.005 | 0.03 |
| TG (18:1/18:2/22:1) | 0.078±0.003 | 0.066±0.005 | 0.032 |

Data are expressed as nmol% of the total lipids; values are expressed as mean ± SE. Data were analyzed by Student's t-test. $p < 0.05$ was considered significant. The table outlines only significant breast milk lipids. N = 40 mother-infant dyads

### 3.5. Influences of Membrane and storage lipids on Atopic disease and growth outcome in babies during the first year of Life

Table 2 shows the significant correlation between membrane lipid and infant atopic disease and growth outcomes. Oleic acid-enriched SM (adjusted correlation coefficient (r)=0.47, p=0.04), TG (r=0.39, p=0.01), DG (r=0.32, p=0.02), and plasmalogen PE (r=0.45-0.61, p<0.05), as well as stearic acid enriched in MG (r=0.37, p=0.02) were positively correlated with the infant FFMI. In contrast, long-chain saturated fatty acids enriched in TG (r=0.38-0.39, p<0.05), SM (r=0.40-0.53, p<0.05), and PI (r=0.49, p=0.01) molecular species were positively correlated with infant FMI. Palmitic and stearic acid-containing phospholipids (r=0.45-0.65, p<0.05) were positively correlated with eczema.

**Table 2:** The association between breast milk lipids and atopic disease or infants’ growth parameters during their first year of life.

| Lipid | Atopic disease or Growth parameter | Correlation |
| --- | --- | --- |
| SM(d16:0/18:1) | FFMI | 0.468 |
| PE (18:0p/18:1) | FFMI | 0.605 |
| PE (16:0p/18:1) | FFMI | 0.450 |
| PE (18:0p/18:2) | FFMI | 0.561 |
| PC (18:0/16:0) | Eczema | 0.537 |
| PS (18:0/20:3) | Eczema | 0.474 |
| PC (16:0/16:0) | Eczema | 0.447 |
| PS (18:0/18:1) | Eczema | 0.652 |
| PE (18:0/16:0) | Eczema | 0.524 |
| SM(d22:0/18:1) | FMI | 0.534 |
| SM(d20:0/18:0) | FMI | 0.397 |
| PI(16:0/16:0) | FMI | 0.486 |
| SM(d20:0/18:0) | Cold | -0.465 |
| TG (18:0e/18:1/18:1) | FFMI | 0.393 |
| TG (18:0e/16:0/18:1) | FFMI | 0.394 |
| MG (18:0) | FFMI | 0.368 |
| DG (15:0/16:1) | FFMI | 0.315 |
| TG (20:0/16:0/18:1) | FMI | 0.376 |
| TG (18:1/18:2/22:1) | FMI | -0.336 |
| TG (18:0/16:1/18:1) | FMI | 0.392 |

Data correspond to significant spearman rank order correlation coefficients. Correlation is significant at $p < 0.05$ . The table shows only significant values: 0.90 to 1.00 (−0.90 to −1.00) - Very high positive (negative) correlation; 0.70 to 0.90 (−0.70 to −0.90) - High positive (negative) correlation; 0.50 to 0.70 (−0.50 to −0.70) - Moderate positive (negative) correlation; 0.30 to 0.50 (−0.30 to −0.50) - Low positive (negative) correlation; 0.00 to 0.30 (0.00 to −0.30) - Negligible correlation. SM-Sphingomyelin, Cer-Ceramide, PC-Phosphatidylcholine, PE-Phosphatidylethanolamine, PS-Phosphatidylserine, PI-Phosphatidylinositol, TG-Triglycerides, MG-Monoglycerides, and DG-Diglycerides.

### 3.6. Diagnostics potential of breast milk functional lipids as Biomarkers in predicting infant development and atopic disease outcomes or phenotype during early life

ROC analysis (Figure 6) showed the r lipid biomarkers associated with infant development and atopic disease outcome. ANCOVA was used to adjust the covariates (Maternal factors). The AUC of ROC output from this study revealed that SM (d16:0/18:1) (AUC-0.788, p<0.0001), PE (18:0p/18:1) (AUC-0.747, p<0.0001), and PE (18:0p/18:2) (AUC-0.717, p<0.0001) can be good biomarkers to predict FFMI in infants during the first year of life: PC (18:0/16:0) (AUC-0.819, p=<0.05), PC (16:0/16:0) (AUC-0.819, p<0.0001), PS (18:0/20:3) (AUC-0.845, p=<0.05), PS (18:0/18:1) (AUC-0.923, p=<0.05), and PE (18:0/16:0) (AUC-0.819, p<0.0001) can be good biomarkers to predict eczema in infants during the first year of life: PI (16:0/16:0) (AUC-0.805, p=<0.05) and SM(d22:0/18:1) (AUC-0.777, p=<0.05) can be good biomarkers to predict FMI in infants during the first year of life.

**Figure 6:**
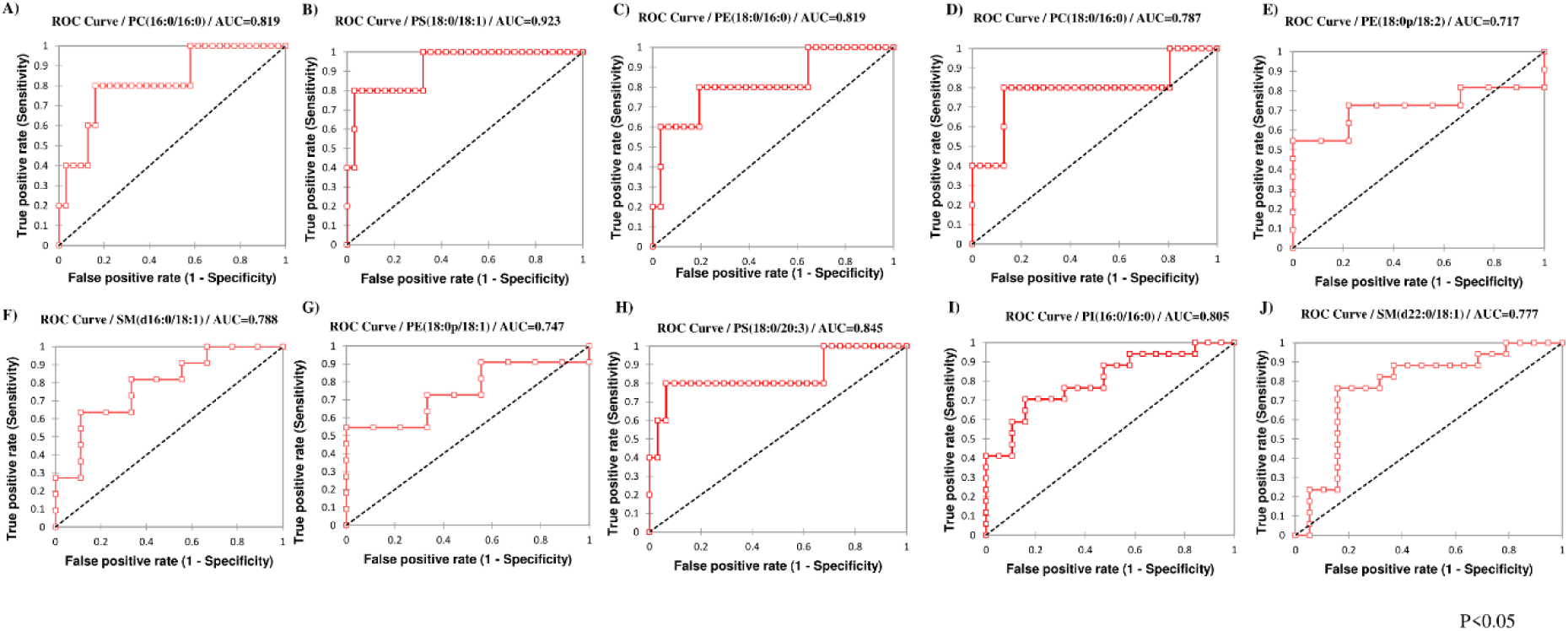
Receiver operator characteristic (ROC) analysis of breast milk lipids significantly associated with infant atopic disease and growth outcome. The area under the curve (AUC) was set to 0.5 and the alpha threshold was set at 0.05.

### 3.7. Lipid network analysis of obese relative to non-obese mother cohorts at one and four-month measurements

The network displays (Figure 7) significant Spearman correlations among lipids (pFDR < 0.05) as edge colors (positive-red; negative-blue). Example: C18:1 shows high fold change lipid, and a positive correlation with TG (18:0/18:1/18:1), TG (18:1/12:0/18:1), and TG (18:0/17:0/18:1). Output from the network analysis demonstrated that SM (d24:0/18:1), DG (14:0/18:2), C(20:3n6), TG (16:0/12:0/14:0), TG (18:0/18:0/20:4), TG (20:0/16:0/18:1), C10, and C14 were significantly increased in obese mothers’ breast milk compared to non-obese mothers’ breast milk. Increases compared to non-obese mothers’ breast milk were observed at one- and four-month post-partum visits. Furthermore, C16, C18, C18:1, PE (18:0p/18:1), PE (18:0P/20:1), TG (18:0/18:1/18:1), SM (d20:1/18:1) were significantly increased in non-obese mothers’ breast milk compared to obese mothers’ breast milk. Increases compared to obese mothers’ breast milk were observed at both one- and four-month post-partum visits. MG (18:0), PC (16:0/18:1), and PE (18:0p/22:6) were significantly increased in non-obese mothers’ breast milk compared to obese mothers’ breast milk in the first month, while PE (18:0P/18:2), TG (4:0/16:0/17:0) decreased at fourth post-partum breast milk.

**Figure 7:**
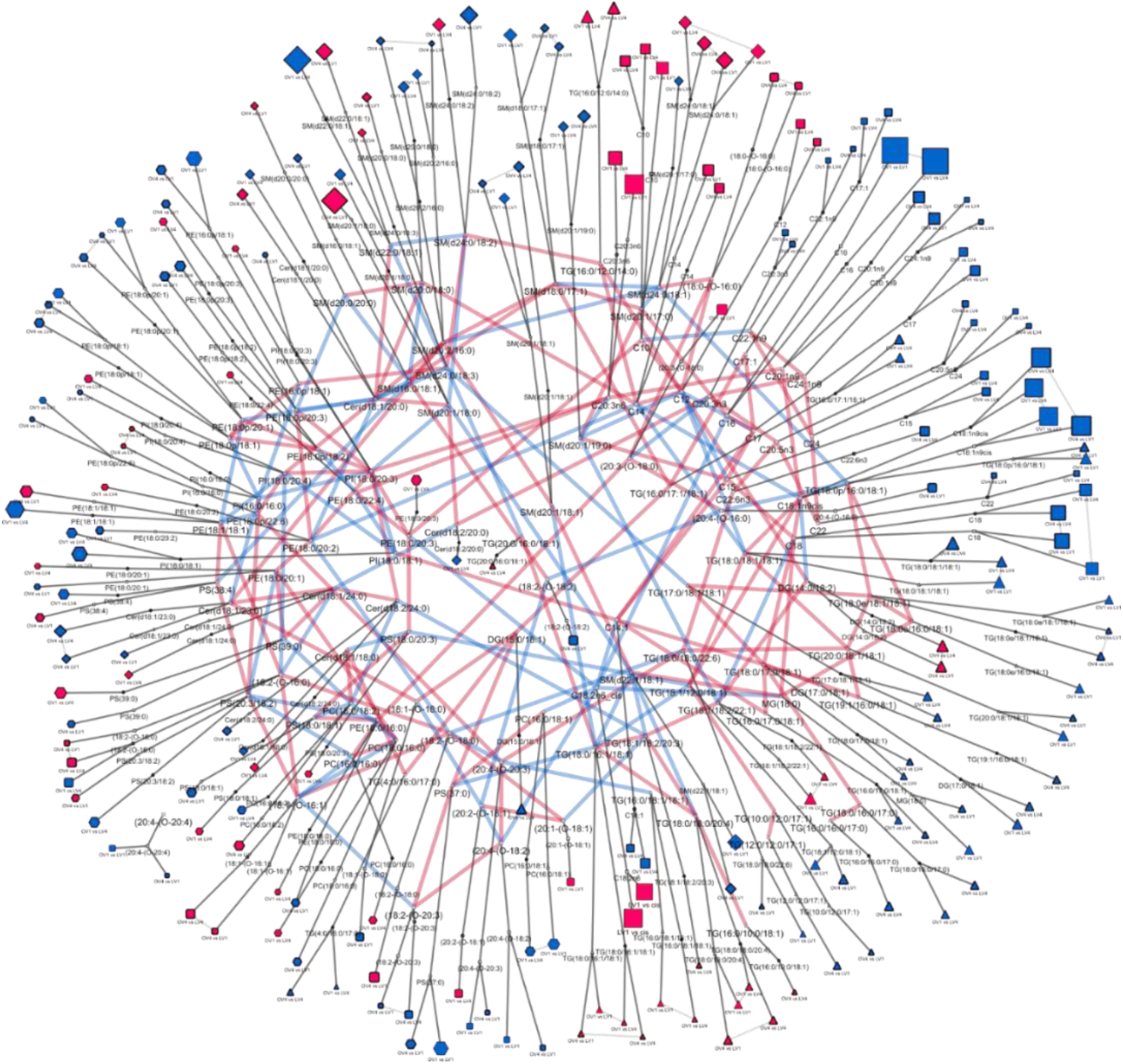
Lipid-lipid partial correlation network displaying differences in obese(O) relative to non-obese(L) cohorts at visit one- and four-month measurements. Network displays significant Spearman correlations among lipids (pFDR < 0.05) as edge colors (positive, red; negative; blue). Statistical contrasts (p-value < 0.05) in obese relative to non-obese cohorts at one and four months are connected to lipid nodes (dark gray edges). Obese vs non-obese comparisons which are not statistically different between time points are connected with light gray edges. The magnitude and direction of fold changes in obese relative to non-obese are shown as node size and color (red, increased; blue, decreased). Comparisons at one and four months are denoted by node border thickness (thin, one month; thick, four months). Lipid classes are denoted with node shape: CER/SM, ceramide, and sphingomyelin; FFA/OxL, free fatty acids, and oxylipins, MAG/DAG.TAG, mono-/di-/triglycerides; PC/PE/PI, phosphatidyl-choline/ethanolamine/serine

### 3.8. Metabolic pathways associated with the altered breast milk lipids

Figure 8 shows the scatter plot of metabolic pathways associated with altered breast milk lipids. These lipids from our data set were predicted to participate in seven metabolic pathways including sphingolipid, glycerophospholipid, linolenic acid, alpha-linolenic acid, arachidonic acid, glycerolipid metabolism, and glycosylphosphatidylinositol-anchor biosynthesis. Sphingolipid metabolism was a relatively high-impact pathway, while glycerophospholipid metabolism was the most significant and impactful pathway. The KEGG pathway was then plotted for further analysis of the high-impact sphingolipid metabolism (Figure 9 (A)) and glycerophospholipid metabolism (Figure 9 (B)) pathways. According to the compound color, the red color means lipid metabolites that significantly influenced the enrichment analysis, and light blue means those metabolites that are not included in the data and are used as a background for enrichment analysis. Furthermore, PE, PC, and PS were important lipid intermediates linking the glycerophospholipid metabolism. However, there were no significant differences in PE, PC, and PS between non-obese and obese mothers’ breast milk. In the sphingolipid metabolism, ceramide was an important lipid metabolite that was significantly higher in obese mothers’ breast milk.

**Figure 8.**
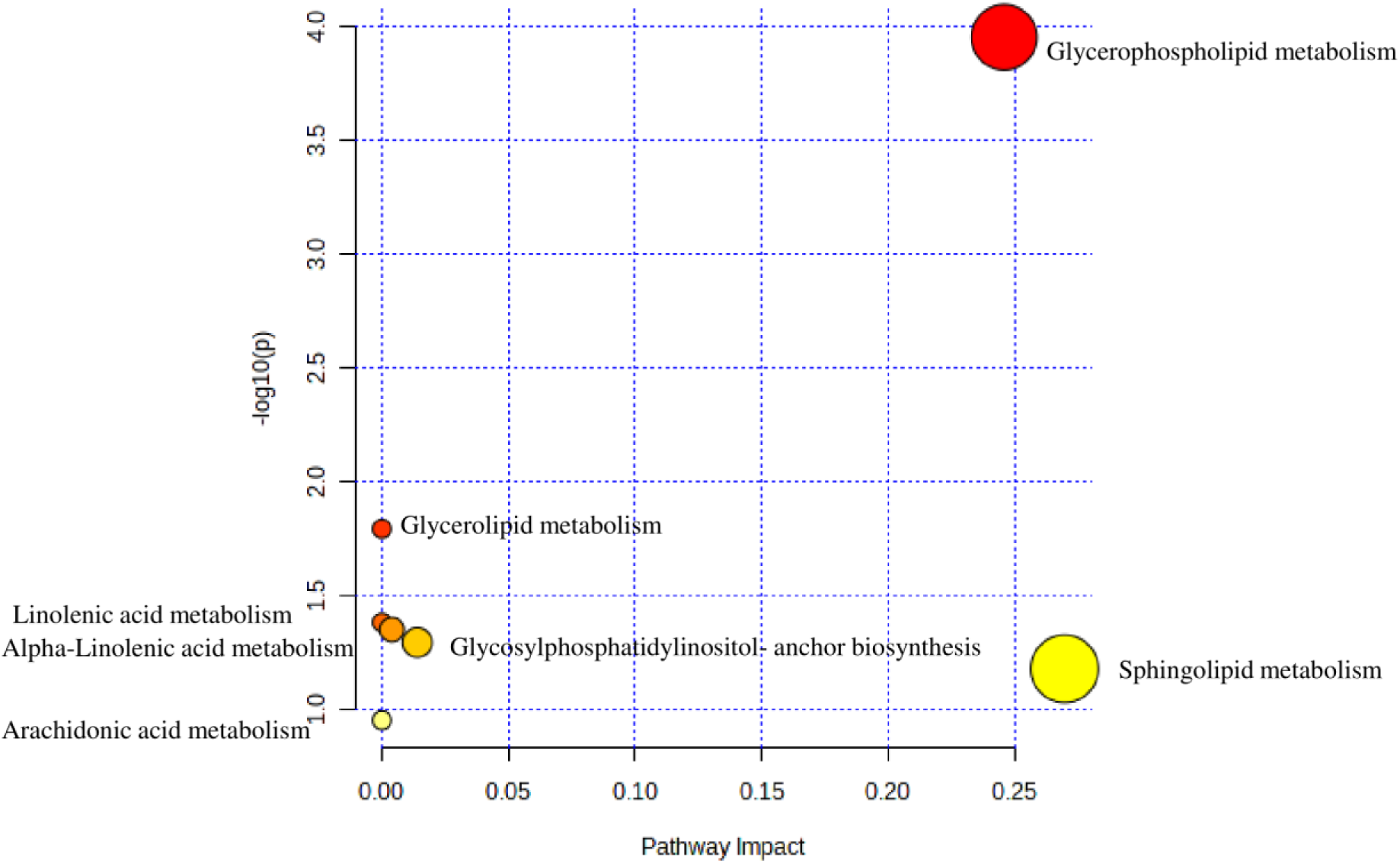
Pathway analysis showing the most important lipid metabolic pathways associated with the altered breast milk lipidome observed between obese and non-obese mothers. Pathway view of statistically significant pathways from the metabolome view based on matched metabolites. The figure shows pathways matched from the lipid metabolites of obese and non-obese breast milk. The pathways are arranged based on the p-value (y-axis), which indicates the pathway enrichment analysis, and pathway impact values (x-axis) representing pathway topology analysis. The node color of each pathway is determined by the p-value (red = lowest p-value and highest statistical significance), and the node radius (size) is based on the pathway impact factor, with the biggest indicating the highest impact.

**Figure 9:**
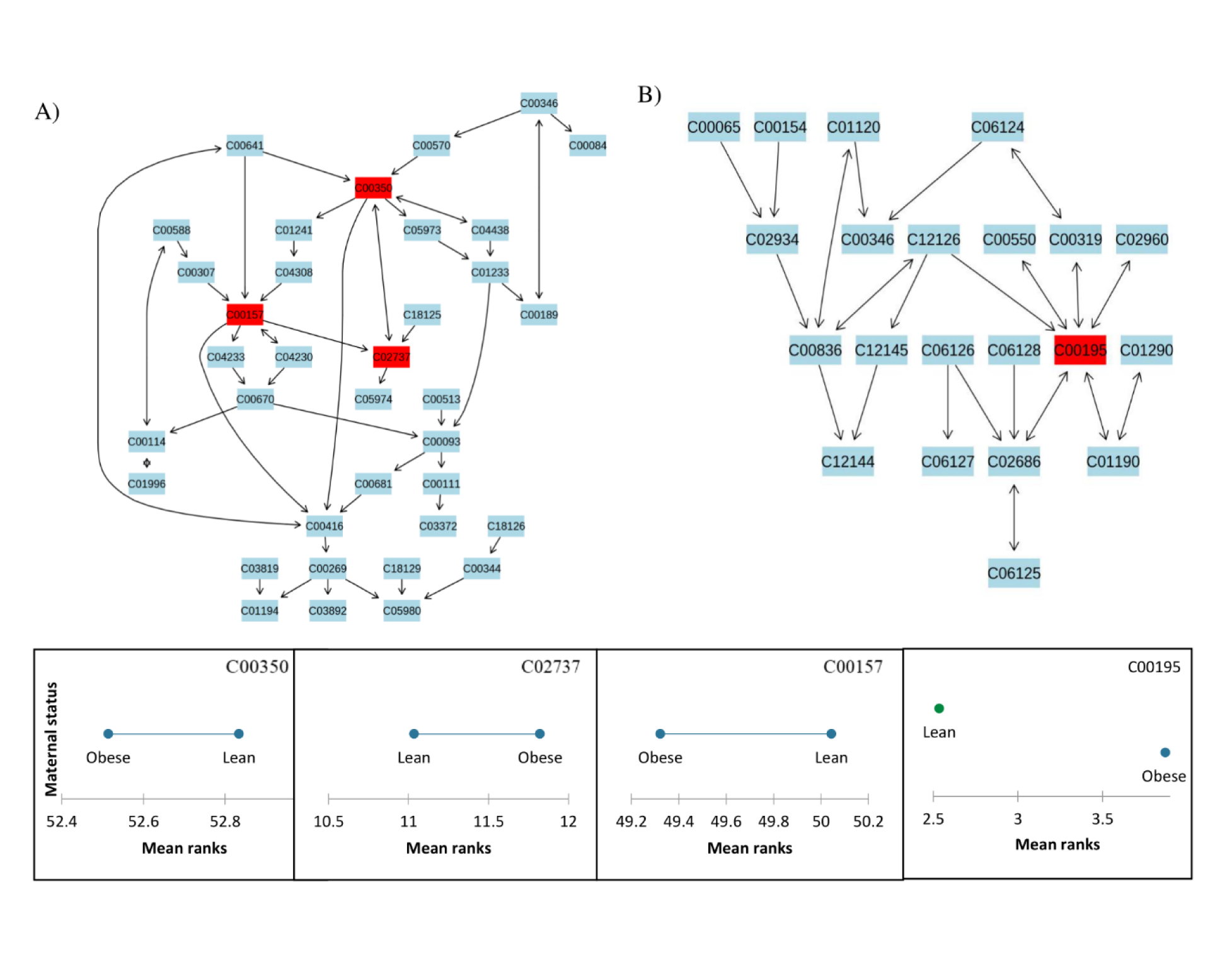
The pathway diagram illustrates the integration and contribution of matched metabolites in glycerophospholipid metabolism (A) and sphingolipid metabolism (B). The metabolites are matched using KEGG identifiers.C00157-Phosphatidylcholine, C00350-Phosphatidylethanolamine, C02737-Phosphatidylserine, and C00195-Ceramide. The demsar plot indicates the mean rank of each metabolite in the obese and non-obese mothers’ breast milk.

## 4. Discussion

Our study investigated i) the associations of maternal body mass index and time postpartum with breast milk lipid composition, ii) how milk lipid composition relates to infant growth, atopic disease, and metabolic outcome; iii) illustrated the benefit of utilizing an advanced lipid bioinformatics approach that is novel in assessing breast milk functional lipidome.

Lipid bioinformatics tools have helped to understand biological phenomena or features through automated data processing using statistical analysis such as pathway and network analysis, as well as lipid modeling to interrogate biological samples such as breast milk ^10,11^ Multivariate analysis methods useful for dimension reduction and group differentiation must be chosen based on the structure of biological data ^22,23^. In our study, RDA was the most appropriate multivariate approach that grouped the lipids based on maternal groups (non-obese and obese) and showed the association between infant atopic diseases and growth outcomes with statistical significance (p-value<0.0001) (Figure 2). RDA describes the linear relationship between multiple dependent and independent variables in the data set ^24^. Additionally, the volcano plot (Figure 3) allowed visualization of the fold-change and significance of lipid concentration in obese and non-obese mothers’ breast milk, identifying lipids characterized by the most discriminating power. Other multivariate analyses such as PCA, PLSDA, and DA were also assessed. However, they were found unsuitable for discerning discreet clusters or features in the breast milk functional lipidome separating the variables under investigation based on the data structure.

Understanding and developing advanced lipid bioinformatics workflow are crucial for assessing the functional breast milk lipidome. For example, the lipid network analysis visualized lipid changes between non-obese and obese cohorts at one and four months and correlations between lipids helped to find the functional relationships between the altered lipids. It revealed the biochemical transformation and structural similarity between lipid species. Biomarkers due to their high predictability, can play an important role in biomarker prediction of possible phenotypic outcomes. The ROC curve evaluates the diagnostic power of a biomarker^25^ while pathway analysis explores unknown functional interactions between lipids and sheds light to find the potential mechanism associated with the altered breast milk metabolism observed between non-obese and obese mothers at one- and four-month post-partum^26^.

Our study found that SM, TG, and PI containing saturated fatty acids, and TG containing long-chain fatty acids were significantly higher in the obese mothers’ breast milk. The SM, DG, and TG containing oleic acid in the sn-2 position TG and MG, containing short chain fatty acids, and plasmalogen containing PE were significantly higher in the non-obese mothers’ breast milk. Lipid network further analysis revealed that C18:0 and C18:1 was significantly higher in non-obese mothers at both post-partum one and four and C12, C14, and C10 were significantly higher in obese mothers’ breast milk, compared to non-obese mothers’ breast milk. These trends with plasmalogen C18:0 and C18:1 levels were confirmed by applying different computational approaches to the same dataset ^7,27^.

Other studies showed that Phospholipids with fatty acids chain lengths shorter than 18 carbon-containing were high in pregnant obese mothers’ serum compared to non-obese pregnant mothers’ serum^28^. The human cross-sectional study showed that the triglyceride concentration was significantly higher in obese mothers’ serum compared to normal-weight mothers’ serum, but they did not show any significant difference in colostrum^29^. Obese mothers showed high levels of palmitic acid, dihomo-gamma-linolenic acid (DGLA), omega 6 polyunsaturated fatty acid, and adrenic acid and low levels of oleic acid and conjugated linoleic acid in their breast milk^27,30^. Maternal factors, such as lactation stage, food intake, and genetic factors influence maternal obesity and breast milk lipid profiles^6,31^. We found that concentrations of phospholipid, sphingomyelin, and storage lipid molecular species were reduced overall over postpartum time. It is supported by other studies that phospholipid and sphingomyelin species were reduced throughout the lactation stages ^32^.

Sphingolipid metabolism and glycerophospholipid metabolism were highly impactful pathways associated with the altered breast milk functional lipidome observed between non-obese and obese mothers based on our study. This is supported by another studies that have reported that breast milk lipids are significantly involved in glycerophospholipid and sphingolipid metabolism ^33^.

## 5. Conclusion

The application of lipid bioinformatics tools enhanced our ability to understand how maternal body mass index influences breast functional milk lipid profile associated with the milk fat globule membrane and the storage center and their effects on infant growth, atopic disease status, and system biology associated metabolomic pathway impacted in early life. By using an advanced lipid bioinformatics approach, we identified key lipid molecular species that were altered in obese and nonobese mothers’ breast milk, connecting these alterations to infant developmental and atopic disease outcomes. Also, our findings showed the importance of lipid metabolomic pathways such as glycerophospholipid and sphingolipid metabolism and their role in infant growth outcomes. Summarizing from a lipid bioinformatics approach, this is one of the first works connecting the functional breast milk lipidome, maternal body mass index of moms, and associations with infant development and atopic disease outcome during the first year of life. However, the study was limited by a smaller sample size. Further work is required with a larger sample size and longer longitudinal follow-up, applying the techniques presented in this paper as well as testing the biomarkers and their ability to predict atopic disease and growth trajectory in infants during early life. Furthermore, these kinds of studies can help determine whether there may be any possible relationships with lactational programming on the trajectory of transgenerational obesity, development, and infant health phenotypes during early or later life.

## Supporting information

C:\Users\Moganatharsa.G\Downloads\New folder (22)

## Author contributions statement

M.G.: writing-original draft. F.E., C.L.W., S.E., C.A., S.S., S.C., and R.T.: conceptualization. S.S., C.L.W and C.A: resources. T.H.P., R.T., M.G.: data curation. M.G.: data analysis. D.G: network analysis. M.G., C.L.W., T.H.P., F.E., S.S., S.C., R.T., and S.E.: writing-review and editing. All authors read and approved the final draft of the manuscript.

## Acknowledgments

Atlantic Canada Opportunities Agency and Memorial University of Newfoundland-Graduate studies.

