## Supplementary material for "Exploring the impacts of human breast milk functional lipidome on infant health and growth outcomes in early life using lipid bioinformatics": C:\Users\Moganatharsa.G\Downloads\New folder (22)

**SUPPLEMENTARY INFORMATION**

**Table S1 Descriptive characteristics of mother–infant dyads**

|  | Non-obese | Obese |
| --- | --- | --- |
| N | 20 | 20 |
| Maternal |  |  |
| Age (years), mean (SD) | 31.87(6.21) | 29.70 (5.97) |
| BMI (Kg/m2), mean (SD) | 25.02(2.49) | 33.92(4.30) |
| Education Level n (%) |  |  |
| Less than high school | 1(5%) | 5(26%) |
| High school | 5(25%) | 2(11%) |
| Some college | 2(10%) | 3(16%) |
| Associate Degree (2-year college) | 1(5%) | 0 |
| Bachelor’s degree | 3(15%) | 5(26%) |
| Master’s degree | 7(35%) | 3(16%) |
| Ph.D./Sc.D./M.D. | 1(5%) | 1(5%) |
| Race/ethnicity n (%) |  |  |
| Asian or Pacific Islander | 2(10%) | 0 |
| Causian | 15(75%) | 9(47.37%) |
| Hispanic | 3(15%) | 6(31.57%) |
| African American | 0 | 4(21.05%) |
| Infant |  |  |
| Birth weight (g), mean(SD) | 3372.55(442.358) | 3242.80(492.776) |
| Gender |  |  |
| Female | 6(31.6%) | 7(36.8%) |
| Male | 13(68.4%) | 12(63.2%) |

*Values were represented as mean ± SD, BMI-Body Mass Index, Education level is the highest level.Abbreviations;SD-Standard deviation ; BMI-Body Mass Index*

| Components | F1 | F2 | F3 | F4 | F5 | F6 | F7 | F8 | F9 | F10 | F11 | F12 | F13 |
| --- | --- | --- | --- | --- | --- | --- | --- | --- | --- | --- | --- | --- | --- |
| Cumulative % of PCA(membranelipid) | 31 | 46 | 56 | 62 | 67 | 71 | 75 | 78 | 80 | 83 | 85 | 87 | 88 |
| Cumulative % of PCA (Glycerolipid) | 21 | 42 | 53 | 60 | 66 | 70 | 73 | 76 | 78 | 80 | 82 | 83 | 85 |

***Table S2 Cumulative percentage of PCA by components of membrane lipid and storage lipids***

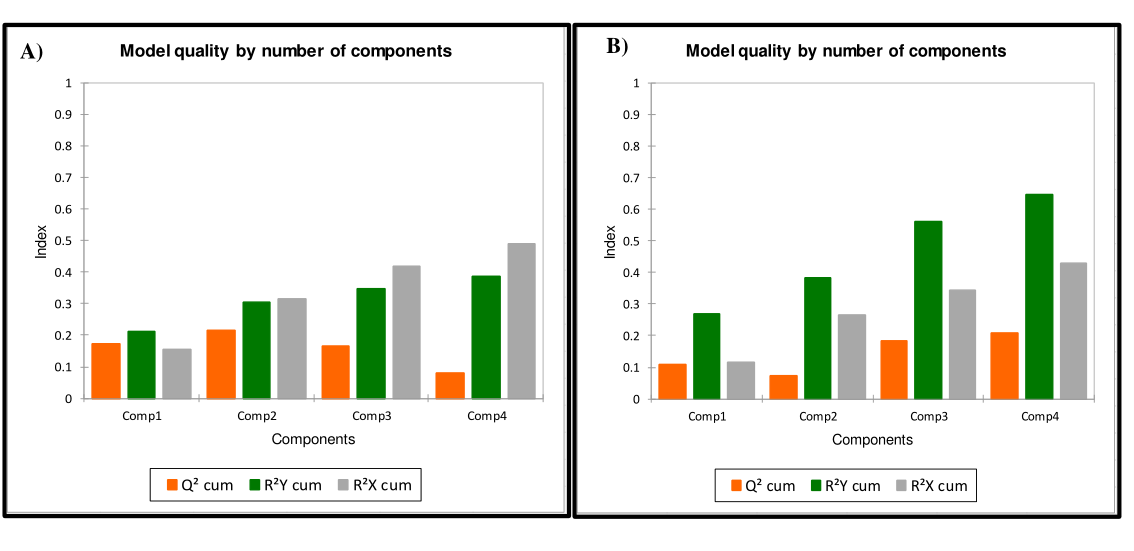
 ***Figure S1 Model quality for partial least squares-discriminant analysis (PLS-DA) of membrane lipids (A) and storage lipids (B)***
